# Reaching the tumor: mobility of polymeric micelles inside an *in vitro* tumor-on-a-chip model with dual ECM

**DOI:** 10.1101/2023.07.14.548885

**Authors:** Alis R. Olea, Alicia Jurado, Gadi Slor, Shahar Tevet, Silvia Pujals, Victor R. De La Rosa, Richard Hoogenboom, Roey J. Amir, Lorenzo Albertazzi

## Abstract

Degradable polymeric micelles are promising drug delivery systems due to their hydrophobic core and responsive design. When applying micellar nanocarriers for tumor delivery, one of the bottlenecks encountered *in vivo* is the tumor tissue barrier – crossing the dense mesh of cells and extracellular matrix (ECM). Sometimes overlooked, the extracellular matrix can trap nanoformulations based on charge, size and hydrophobicity. Here we used a simple design of a microfluidic chip with two types of ECM and MCF7 spheroids to allow “high throughput” screening of the interactions between biological interfaces and polymeric micelles. To demostrarte the applicapility of the chip, a small library of fluorescently labelled polymeric micelles varying in their hydrophilic shell and hydrophobic core forming blocks was studied. Three widely used hydrophilic shells were tested and compared, poly(ethylene glycol), poly(2-ethyl-2-oxazoline), and poly(acrylic acid), along with two enzyamticaly degradable dendritic hydrophobic cores (based on Hexyl or Nonyl end-gorups). Using ratiometric imaging of unimer:micelle fluorescence and FRAP inside the chip model, we obtained the local assembly state and dynamics inside the chip. Notably, we observed different micelle behaviors in the basal lamina ECM, from avoidance of ECM structure to binding of the poly(acrylic acid) formulations. Binding to basal lamina correlated with higher uptake into MCF7 spheroids. Overall, we proposed a simple microfluidic chip containing dual ECM and spheroids for the assessment of the interactions of polymeric nanocarrires with biological interfaces and evaluating nanoformulations’ capacity to cross the tumor tissue barrier.

## Introduction

In the last seven decades, nanoparticles received increasing attention as possible vehicles to transport drugs in a targeted manner, and serve as drug delivery systems (DDSs) for cancer treatment.^1^ Their use promises to alleviate side effects caused by systemic administration and to increase the therapy effectiveness. Despite the intensive research, only a handful of DDSs reached the clinic. One of the reasons that makes DDS design very challenging is the lack of comprehensive testing platforms. While *in vivo* experiments using animal models are widely used for mimicking the tumor environment, differences to the human counterparts, the complex protocols as well as ethical issues make the emerging 3D *in vitro* platforms a more attractive tool to mimic the interactions inside the human body and predict the efficacy of the studied DDSs.^2^ Once a DDS enters the human body via intravenous injection, there are several bottlenecks it needs to surpass.^3^ In the blood circulation, DDS encounters sudden dilution, sheer stress and blood proteins, which can interact by creating a protein corona, leading to clearance by the spleen or kidneys. Then, DDS needs to extravasate at the target site, reaching the tissue barrier: extracellular matrix filtration, high intratumoral pressure, passage through several layers of cells; finally reaching the target cells, where DDS should release the cargo to perform its intracellular activity.^4^ Although much attention was given to the circulation and cellular internalization steps, there is less focus on the interaction with the tumor tissue barrier. After extravasation, the nanoparticles would reach dense layers of cells and extracellular matrix (ECM).^5^ From a structural view, ECM is a mesh that gives the shape to organs and tissues and directs cellular movement. Yet, from a functional view, recent studies have shown that ECM can filter charged nanoparticles of either sign (positive or negative).^6,7^ It can also trap large particles^8^ or cause destabilization through hydrophobic interactions (which can cause DDSs to lose their cargo).^5^ Also, the high intratumoral pressure present in some cases can deter the entry of DDSs. It is known that cancer tissues have different, much stiffer microenvironments compared to those of normal tissues, and different matrix-associated protein composition. These features affect cell differentiation, proliferation and migration, as well as gene expression and response to anticancer drugs, contributing to the tumorigenic microenvironment.^9^ Often, the presence of ECM is ignored when testing anti-cancer formulations. Most 2D cell cultures lack a viable ECM, while animal models can have a different ECM compared to the human environment. However, ECM density was shown to directly affect tumor penetration for different sizes of polymeric micelles *in vivo*.^8^ Thus, being able to test the passage of DDSs through tumor ECM would be one more step of the puzzle in aiding the design of effective DDSs. This could be achieved using an *in vitro* platform mimicking the tumor ECM in which the interaction with different nanoformulations could be tested. What should such a platform contain? Extracellular matrices around the body have various compositions. Thus, for designing a comprehensive test platform, there are several ECM types that should be taken into account. One of the most important is the basal lamina, which has proven nanoparticle filtration properties^6^. Basal lamina is a thin ECM layer that creates the inner lining of many epithelial, muscle and endothelial tissues, including the walls of blood vessels. Basal lamina is an intertwined mesh of laminin, collagen type IV and heparan sulphate chains, crosslinked by several connecting molecules.^10^ The reticular mesh-like structure of basal lamina is very different from the ECM typically found inside tumors. For instance, the desmoplastic tumors such as breast cancer, pancreatic or prostate cancer have an increased deposition of macromolecules otherwise specific to wound healing sites^11^, resulting in the disorganized ECM of a wound that does not heal. Such macromolecules include collagens type I, III and V (which have a fibrillary structure) and hyaluronic acid (especially ones with low molecular weight, which promote inflammation,^12^ but also high molecular weight segments which contribute to cancer resistance^13^). These macromolecules create a high local pressure due to water retention, greatly affecting the flow of nutrients and signaling molecules in the area^14^.

For reconstituting the basal lamina ECM and the intra-tumoral ECM *in vitro*, widely used models are Matrigel – basement membrane extract from Engel-breth-Holm-Swarm murine sarcoma cells^15,16^; and collagen type I, respectively.^17,18^ Both are very well researched for their stiffness and microarchitecture formation to correspond to the tumor environment. Moreover, Matrigel is able to exhibit nanoparticle filtration effects based on NPs surface charge, while a simple mix of the basal lamina components cannot recapitulate this feature.^6^ Notably, both positively and negatively charged NPs can be retained. Furthermore, size filtration and hydrophobic interactions^5^ should be also taken into account. Therefore, we introduce here a simple testing platform that can be obtained by combining both types of ECM models, representing the ECM barriers of blood vessels and intra-tumoral ECM, together with MCF7 breast cancer cellular spheroids, for an easy screening of DDS formulations in their ability to cross the tumor tissue barrier.

To demonstrate the applicability of this model, we will use it to study the mobility and targeting of polymeric micelles, which are a promising category of DDS thanks to their modular design and the capacity to encapsulate hydrophobic drugs. A small library of fluorescently labelled micelles was formulated varying in their hydrophilic shell, while maintaining their hydrophobic block.^19^ This is an important structural feature as it allows us to study the outside shell, which is the first to interact with the biological interfaces and its charge, hydrophilicity and organization deeply influence the material-cell interactions. Poly(ethylene glycol) (PEG) is the most commonly used hydrophilic polymer but its wide use is already causing an immune response in the population. One-to-one comparison was made between the effect of three common hydrophilic shells PEG, poly(2-ethyl-2-oxazoline) (PEtOx), and poly(acrylic acid) (PAA), on the interactions of polymeric micelles inside a 3D microfluidic chip with a dual-ECM model of breast cancer and MCF7 spheroids. Notably, taking advantage of their fluorescent labelling, we measured both the assembly state of the micelles and the diffusion inside ECM at the same time, by using ratiometric imaging and fluorescence recovery after photobleaching (FRAP). We observed differences in the interaction behavior of different micelles with ECM components, especially for the basal lamina model.

## Materials & Methods

### Chip preparation

The commercial microfluidic chip DAX-1 (AIM Biotech) was filled in the middle channel with ECM from Engelbreth-Holm-Swarm murine sarcoma cells (Sigma Aldrich), 2x diluted in full DMEM (10% FBS, from Thermo Fisher) to a final protein concentration of 5.25 mg/mL. The ECM was allowed to gelate inside the chip for 15-20 min at 37° C, 5% CO_2_, placed on two PDMS supports of 4 mm thickness, one at each end of the chip, inside a Para-film-sealed Petri Dish. After ECM gelation, the right side channel was filled with collagen-hyaluronic acid mix containing MCF7 spheroids. The mix preparation was adapted from AIM Biotech general protocol v.5.3. Collagen gel was prepared at a final concentration of 2.5 mg/mL gelated inside the chip at pH 7.4 and 37° C, conditions which mimic tumor environment^20^. Briefly, collagen type I from rat tail acid solution (Corning) was brought to pH 7.4 on ice by pre-mixing with Phenol Red and PBS 10x (both from Sigma Aldrich) as 10% of final volume, then adding sodium hydroxyde 0.5 N (PanReac NaOH pellets dissolved in MilliQ water) until the color changed to faint pink. Hyaluronic acid sodium salt from rooster comb (Sigma Aldrich) dissolved in MilliQ water was added to a final concentration of 0.8 mg/mL. The gel mix was used to resuspend pre-made MCF7 spheroids after 5 min centrifugation at 200 g. MCF7 spheroids were obtained by seeding 1000 cells/well in a 96-well low-attachment NUNC Sphera plate, 48h prior to chip preparation. The right side-channel of the chip was filled with the final mix, then the chip was incubated at 37° C, 5% CO_2_ for 30 min using the same Parafilm-sealed Petri Dish setup. After gelation, the left side channel was filled with micelle solution of 160 μM, diluted 3x in full DMEM (from 480μM in PBS, pH 7.4). Micelle solution was then added to the reservoirs on the left side of the chip and allowed to diffuse inside the chip overnight at 37° C, 5% CO_2_.

### Cell culture reagents

MCF7 cells were cultured in full DMEM medium (Dulbecco’s modified Eagle Medium 1x with added 4.5 g/L D-glucose, L-glutamine and pyruvate, from Thermo Fisher), supplemented with 10% fetal bovine serum, heat inactivated (Gibco, Thermo Fisher) and 1% penicillin/streptomycin (Biowest). Trypsin 25% EDTA (Thermo Fisher) was used for cell detachment.

### Gel labeling

ECM from Engelbreth-Holm-Swarm murine sarcoma cells, 2x diluted in full DMEM (10% FBS) to 5.25 mg protein/mL was mixed with Cyanine3 NHS ester (Lumiprobe, dissolved in DMSO) to a final dye concentration of 19 μM and allowed to react for 1h on ice. Then, the reaction mix was added inside the chip middle channel and allowed to gelate for 15 min at 37° C, 5% CO_2_ in a Parafilm sealed Petri Dish. Rat tail collagen type I (at final concentration 2.5 mg/mL and pH 7.4) and hyaluronic acid (0.8 mg/mL) mix was reacted with 10 μM Cyanine5 NHS ester (Lumiprobe, dissolved in DMSO) for 45 min on ice, then added to the right side channel of the chip after the gelation of the middle channel. The collagen-HA mix was allowed to gelate for 30 min at 37° C, 5% CO_2_. The remaining side-channel of the chip was filled with PBS (pH 7.4) and the chip was imaged immediately in Zeiss LSM 800 confocal microscope, using 561 nm and 640 nm excitation for Cy3 and Cy5 channels respectively.

### Live/dead assay

A chip was prepared containing MCF7 spheroids and allowed to grow for 24h in full DMEM. Using only the flow channel, the chip was first washed with serum-free DMEM, which was then used for all further steps. Then, calcein AM (Sigma Aldrich) solution 10 μM in serum-free DMEM was added to the left side channel of the chip and allowed to distribute inside the chip for 1h at at 37° C, 5% CO_2_. Then, the solution was replaced by propidium iodide in serum-free DMEM, for 10 min. A final wash was performed, leaving the chip with serum-free DMEM during imaging in confocal microscope.

### Micelle characterization (Zeta potential and DLS)

Previously reported amphiphiles^19^ were used to prepare micelle solution of 80 μM in PBS (pH 7.4). A Malvern Zetasizer Nano ZS was used for ζ-potential measurements in plastic cuvettes and dynamic light scattering in low-volume quartz cuvettes for determining the hydrodynamic size. Measurements were performed in triplicate.

### Confocal microscopy

Imaging was performed on a Zeiss LSM 800 confocal laser scanning microscope equipped with two PTM Multi Alkali detectors, using the Zen 2.3 (blue) software. Images were acquired with a plan apochromat 20x / 0.8 M27 objective and using 37° C, CO_2_/O_2_ incubation. A diode laser 405 nm (5 mW) at 1% power was used for excitation, while the emission was collected in two different channels: 400-500 nm for unimers and 500-700 nm for micelle fluorescence respectively. The two channels were summed in Fiji ImageJ^21^ to obtain the “total fluorescence” images; the unimer signal was divided by the micelle signal after background removal to obtain “ratiometric” images.

### FRAP

Fluorescence recovery after photobleaching was performed in the same LSM 800 confocal microscope using the 20x objective and the dual channel acquisition. A circular region of 35.3 μm in diameter was used for bleaching, in the center of a 103×103 μm image (zoom 3.1x, 256×256 pixels, 16 bit, unidirectional). A total of 4 minutes experimental time were recorded with 102.4 ms/frame and 405 nm excitation at 1%, including 10 frames pre-bleach and the bleaching time of 3,14 sec localized as 10 iterations with 100% laser power inside the bleach area. An overview image of the area (638.9 μm square, zoom 0.5x) was captured before each FRAP measurement. For control measurements, the same micelle solution in full DMEM at 160 μM was added onto a glass slide with two layers of double-sided sticky tape and coverslip on top, then sealed with nail polish to prevent drying. All FRAP measurements were performed with 37° C and CO_2_/O_2_ incubation.

Post-processing of FRAP data was done using Fiji ImageJ^21^ and easyFRAP^22,23^. The fluorescence inside the bleached area was measured using a smaller circle (30 μm in diameter) to avoid measuring any subtle drift. A double normalization of the recovery data was performed with the online easyFRAP tool, correcting for photobleaching during acquisition by using the mean fluorescence of the whole image. Also, we used the assumption that the first post-bleach value in the bleach region is the “background” value. Then, one-component exponential curve fitting was performed in GraphPad Prism (GraphPad Software, San Diego, California, USA), on the normalized data of all repeated measurements, with “Y_0_” constrained to “0”, after removing the pre-bleach values. Taking the half-time obtained in the fitting, we calculated the diffusion constants using the simplified equation of Soumpasis et al.^24^, which assumes instantaneous bleach: D = 0.224 · r_n_^2^/ τ_1/2_, where r_n_ is the nominal radius of the bleach area and τ_1/2_ is the half-time of fluorescence recovery, while 0.224 is a coefficient numerically determined for aqueous environment.

## Results & Discussion

### Experimental setup

In order to model the tumor ECM barrier in a simple *in vitro* testing platform, we used two ECM types and breast cancer MCF7 spheroids inside a microfluidic device. Notably, we intended to mimic the tissue barrier without the extravasation step through the endothelial layer which we addressed in a previous study^25^. As the base for the microfluidic device, we used the commercially available DAX-1 microfluidic chip model from AIM Biotech. DAX-1 has several advantages, especially the ease of reproducibility, being optically transparent (unlike other chips made of PDMS) and having a bottom permeable to CO_2_/O_2_, making it compatible with live cell culture. Also, the platform is versatile enough to implement our idea of dual ECM, as the chip has three microfluidic channels separated by triangular pillars, which allow filling the channels with separate types of gel (Figure 1A). The middle channel is 1.3 mm wide, while both side channels are 0.5 mm wide, with a height of 0.25 mm and 10.5 mm channel length. We chose to fill the middle channel with a gel model for basal lamina: ECM from Engelbreth-Holm-Swarm murine sarcoma cells. After gelation of the middle channel, we filled one of the side-channels with a gel model for desmoplastic tumors, a mix of rat tail collagen type I and hyaluronic acid. In this “tumor ECM” gel we embedded MCF7 spheroids as a widely used model for breast cancer (Figure 1B). In order to obtain a collagen gel microarchitecture resembling solid tumors, we performed the gelation of the collagen mix using a final collagen concentration of 2.5 mg/mL, pH 7.4 and 37° C^20^. After gelation, we used the remaining side-channel for adding the solution of polymeric micelles in cell media (full DMEM, with 10% FBS). After equilibration time overnight, we performed two types of measurements in a confocal microscope: ratiometric imaging for determining local assembly state of the micelles and fluorescence recovery after photobleaching (FRAP) for measuring unimer and micelle dynamics in different parts of the chip (Figure 1).

**Figure 1.**
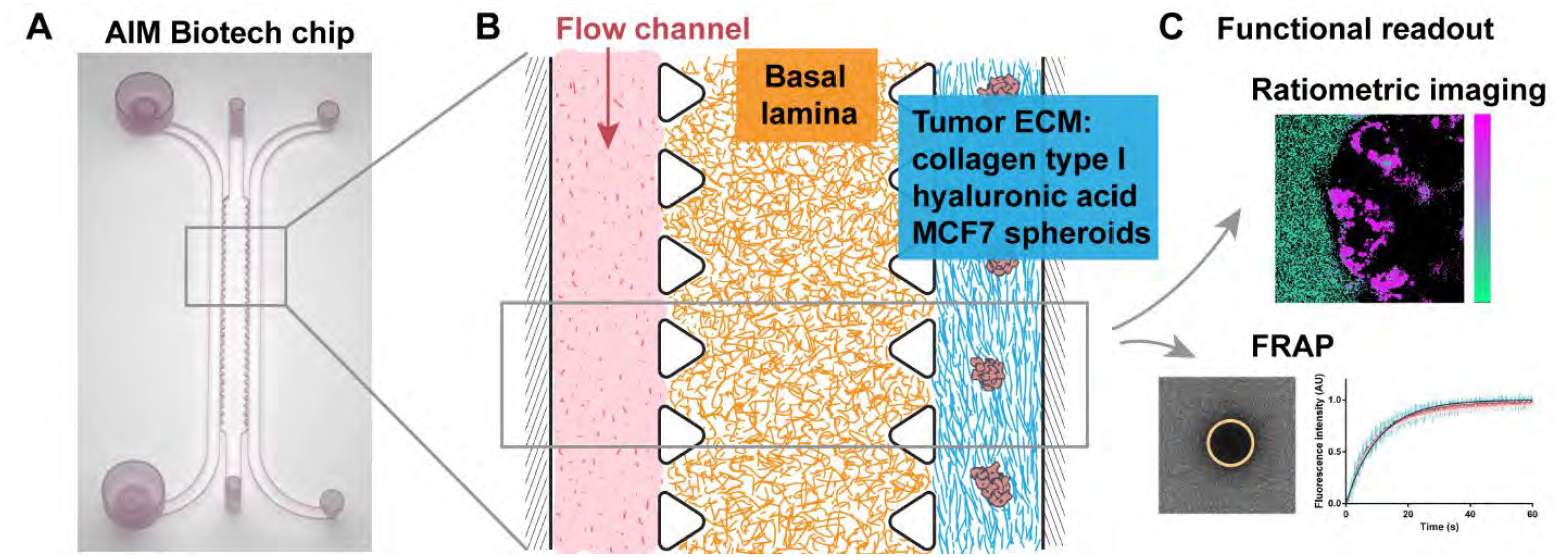
: Experimental setup. A commercial microfluidic chip from AIM Biotech, with three channels separated by triangular pillars (A) is filled in the middle channel with a model of basal lamina ECM from Engelbreth-Holm-Swarm murine sarcoma cells at 5.25 mg/mL and in the right side-channel with a gel mix of collagen type I (2.5 mg/mL, pH 7.4) and hyaluronic acid (0.8 mg/mL), representing the tumor ECM, in which are embedded spheroids of MCF7 breast cancer cell line (B). The micelle sample is added to the flow channel as a 160μM solution in full DMEM (10% FBS) and allowed to diffuse for 24h before doing a functional readout in the confocal microscope, either as ratiometric imaging or as fluorescence recovery after photobleaching (FRAP) in different locations inside the chip (C).

### Chip validation

Before testing micelle’s interactions in the dual-ECM chip, we assessed the integrity of the proposed 3D model. In order to validate if the two types of ECM gels are located in the correct compartment after dual-step filling, we pre-labeled each gel mix with either Cy3 or Cy5 dyes using EDC-NHS reaction. This way, all the components of the basal lamina gel and “tumor ECM” mix should be visible. A transversal view inside the gel-filled chip revealed that the two gel types remained in the expected chip compartments, in the middle channel and side-channel respectively, without mixing (Figure 2A, B).

**Figure 2.**
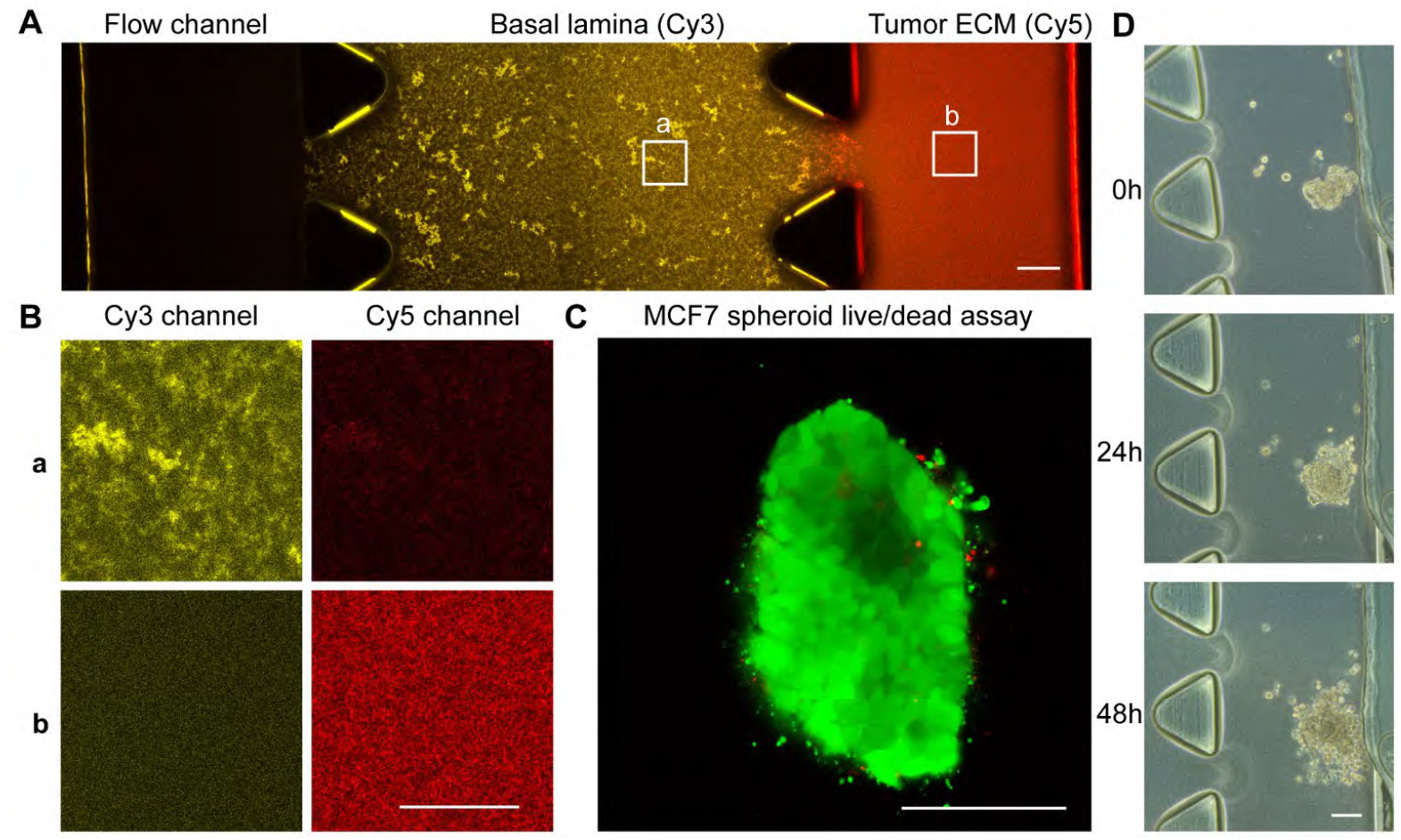
: Validation of ECM distribution and spheroid viability inside the chip. (A) Overview of ECM distribution inside the chip using Cy3-labeled basal lamina gel (ECM from Engelbreth-Holm-Swarm murine sarcoma cells) and Cy5-labeled tumor ECM model (mix of collagen type I and hyaluronic acid), labeled using EDC/NHS reaction. (B) Zoom into (a) Basal lamina and (b) tumor ECM, showing the fluorescence in Cy3 (yellow) and Cy5 (red) channels. (C) Live/dead assay of MCF7 spheroid inside tumor ECM channel, after 24h inside chip, stained with calcein (green) and propidium iodide (red) (image shown after log transformation). (D) MCF7 spheroid growth inside tumor ECM channel. Scale bar is 100 μm for A, C, D and 50 μm for B.

Another step to validate the chip model was to assess spheroid viability and growth. We used a live/dead assay with calcein and propidium iodide in order to visualize the viable and dead MCF7 cells, respectively, inside the chip (Figure 2C). As most of the signal comes from calcein, we concluded that most cells remained viable. This is supported also by observing the spheroid growth from 0 to 48h inside the chip (Figure 2D). Based on the growth images, we decided to use the 24h time point for micelle measurements in the chip, since at 48h the spheroids seem to lose the round shape.

### Micelles characterization

The dual-ECM microfluidic chip model allowed us to compare the penetration capacity of different micelle formulations into a relevant model of tumor extracellular environment, while also comparing the micelle’s internalization capacity into 3D spheroids. Figure 3 presents a graphical overview of the molecular design of the polymeric amphiphiles that were used to prepare the micelles investigated in this study. Three widely known polymers: PEG, PEtOx and PAA, with similar molecular weights were used as the shell forming hydrophilic blocks. The hydrophilic polymers were clicked together with a hydrophobic dendron with four esterase-cleavable chains of either 6 (“Hex”) or 9 carbons (“Non”) in length (Figure 3). Their synthesis was previously described in detail.^19^

**Figure 3.**
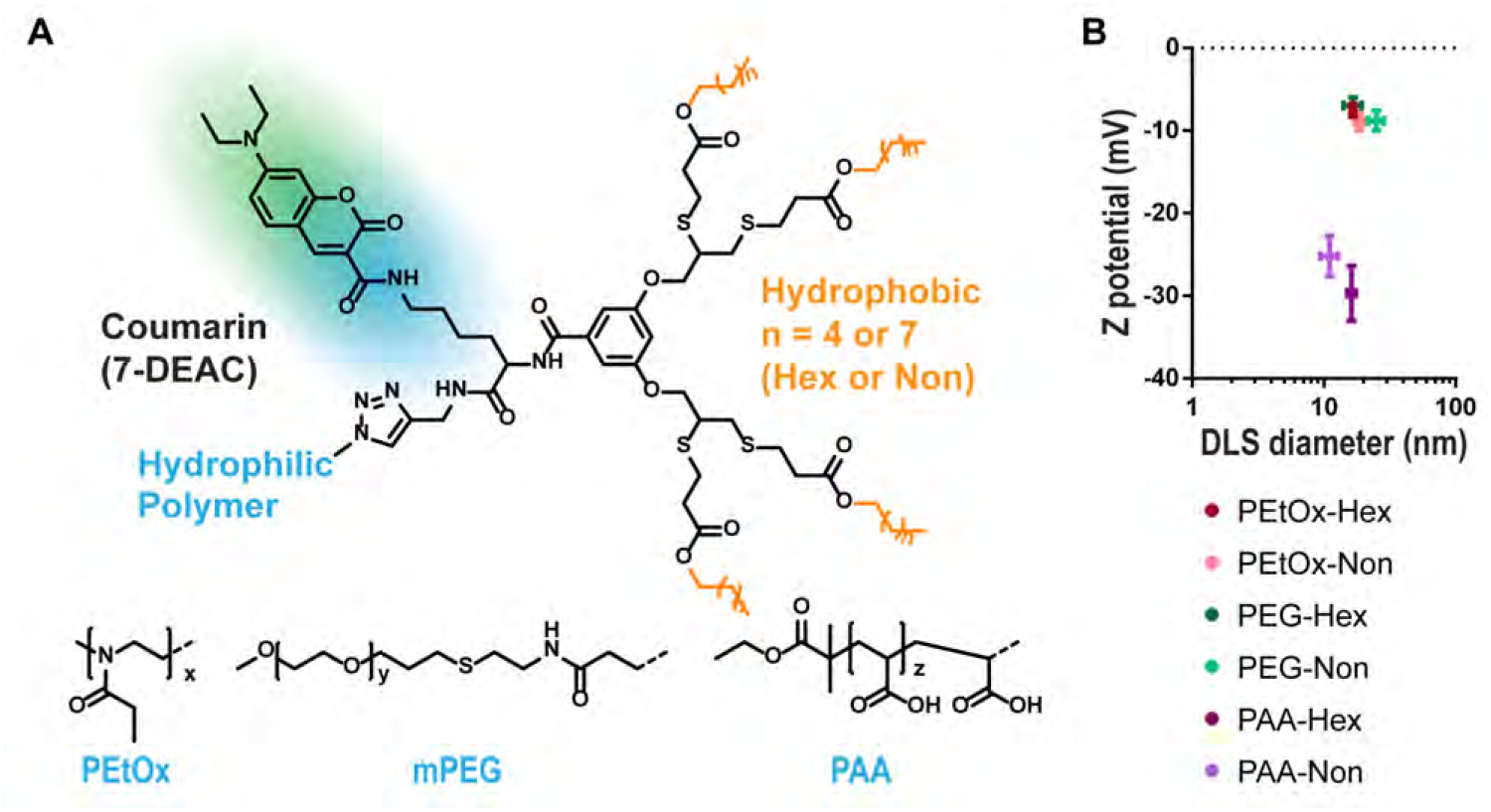
: Micelles structure and characterization. Chemical structure of amphiphiles formulations with 3 different hydrophilic groups (PEtOx, PEG or PAA) and 2 lengths of the hydrophobic ends (“Hex” or “Non”), labeled with 7-DEAC (A). Zeta potential measurements plotted versus hydrodynamic size by DLS of the 6 micelle formulations in PBS, pH 7.4, are shown ([amphiphile] = 80μM) (B).

Additional important aspect in our choice of micelle design is the fluorescent label based on 7(diethylamino)coumarin-3-carboxylic acid (7-DEAC) which changes its emission spectra due to excimer formation when dyes are in close proximity inside the micelle^26,27^. The emission peak thus reflects the assembly state of the polymeric amphiphiles: 480 nm for unimer form and ∼540 nm for micelle. This allows to visualize if micelles disassemble at different locations inside the chip. Overall, the well-defined structure and the fluorescence reporting mechanism gave us the advantage of distinguishing between the effects of the different hydrophilic shells and those that result from the hydrophobicity of the micellar core.

The micelles’ size and charge were assessed by dynamic light scattering (DLS) and Zeta potential measurements using a micelle solution obtained through self-assembly in PBS (pH 7.4) at amphiphiles concentration of 80 μM. DLS measurements showed that micelle size is in the range of 10-30 nm (Table 1), which were similar to our previous studies^19^. Based on their dimensions, we would not expect them to be trapped inside the ECM gels, which under current conditions would have the pore size of a few μm^28^. Instead, interactions with ECM would be due to charge or hydrophobicity. For assessing micelle surface charge, we performed Zeta potential measurements. As expected, PEtOx and PEG micelles showed rather neutral surface charges (−7 to -9 mV in PBS), while PAA polymers had a negative surface charge, with an average of -29 mV for PAA-Hex and -25 mV for PAA-Non (Figure 3B). Overall, the size and charge of the micelles are consistent with our previous study, allowing us to assess their interactions with the different types of ECM in our chip model.

**Table 1.** DLS and Z potential measurements of micelle solution in PBS pH 7.4, ([amphiphile] = 80 μM).

|  | PEtOx-Hex |  | PEtOx-Non |  | PEG-Hex |  | PEG-Non |  | PAA-Hex |  | PAA-Non |  |
| --- | --- | --- | --- | --- | --- | --- | --- | --- | --- | --- | --- | --- |
| DLS (nm) | Mean | SD | Mean | SD | Mean | SD | Mean | SD | Mean | SD | Mean | SD |
|  | 17 | 0.95 | 18 | 1.3 | 17 | 2.6 | 25 | 3.47 | 16 | 1.19 | 11 | 1.66 |
| Z potential (mV) | Mean | SD | Mean | SD | Mean | SD | Mean | SD | Mean | SD | Mean | SD |
|  | -8 | 0.86 | -9 | 1.01 | -7 | 0.98 | -9 | 1.2 | -30 | 3.31 | -25 | 2.48 |

### Confocal imaging

Once the characterization of the micelles in solution was completed and micelles were confirmed to be non-toxic to MCF7 cells in a Presto Blue cytotoxicity assay (Supplementary Figure 1), we moved on to testing micelle distribution and dynamics inside the chip. We filled the “flow channel” with micelle solution ([amphiphile] = 160 μM in full DMEM, 10% FBS) and allowed micelles to distribute inside the chip via passive diffusion during 24h incubation at 37° C, 5% CO_2_. In this case, we avoided the use of a microfluidic pump, since the intention was to mimic the diffusion of nanocarriers inside tumor tissue after the extravasation step. Using a confocal microscope we imaged the coumarin-labeled amphiphiles inside different parts of the chip with 405 nm excitation. Acquisition was split into two channels that reflect the coumarin spectral shift between unimer and micellar states: the unimers channel from 400 to 500 nm and the micelles channel from 500 to 700 nm. As a post-processing step, we summed the two channels to create a “Total fluorescence” image in order to compare overall distribution and intensity. Also, we divided the unimer channel by the micelle channel to obtain a “Ratiometric” image, which indicates the spatial distribution of unimers and micelles in different chip locations.

Looking at the total fluorescence images inside the basal lamina (middle channel of the chip) (Figure 4A top row, 4B; Supplementary figure 2), we observed that the amphiphiles PEtOx-Non and PEG-Non, which have a more hydrophobic block and hence from more stable micelles, showed a similar “dark” structure of ECM (which was not labeled), meaning that these micelles were basically avoiding the basal lamina. In contrast, the negatively charged polymers PAA-Hex and PAA-Non showed a “bright” structure of the ECM, which suggests that they bind to the basal lamina mesh and accumulate on it. The local accumulation of PAA polymers onto the basal lamina is also supported by an overall higher fluorescence intensity compared to other chip compartments (Figure 4D). The other two formulations, PEtOx-Hex and PEG-Hex were found to be in between their more hydrophobic Non-based analogues and the PAA based polymers, not showing a repulsion, but a rather homogenous distribution of fluorescent signal with occasional brighter spots (Figure 4A, B; Supplementary figure 2). In this case, we can assume there is a small degree of interaction with the basal lamina, but not to the point of a visible accumulation on the mesh structure. This would mean there is more interaction with the ECM for Hexyl compared to Nonyl formulations, probably due to the less stable, and consequently, more dynamic nature of the Hexyl micelles, due to the lower hydrophobicity of the core forming dendrons. Notably, the small size of the micelles, 15-25 nm in diameter, and free unimers (expected to be 5-10 nm) is significantly smaller than the expected pore size of the reconstituted basal lamina mesh (∼2 μm).^6^ Thus, we attribute the observed accumulation to the interactions of the polymers with the ECM interface and not due to their size.

**Figure 4.**
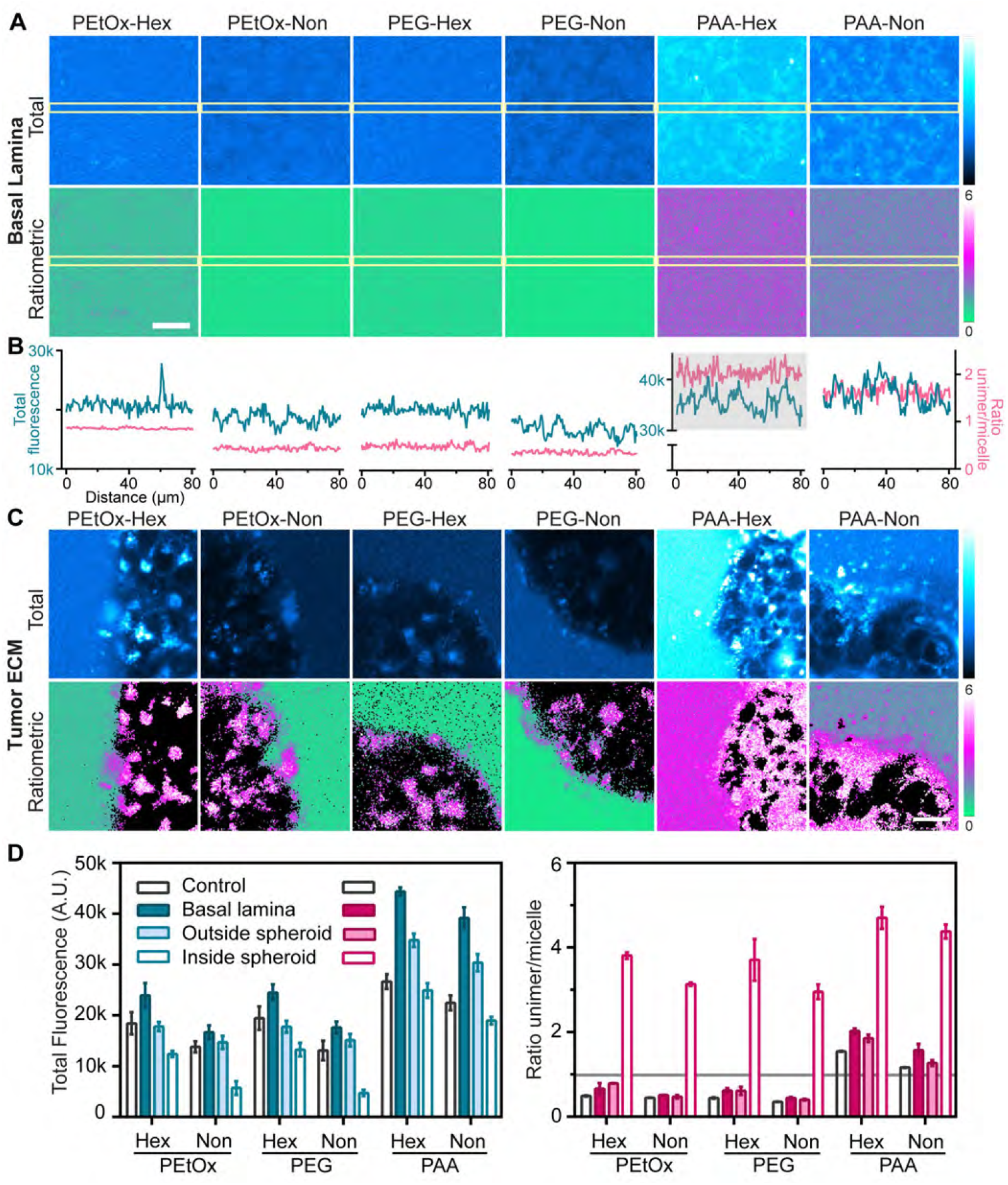
: Confocal imaging of micelles inside different chip compartments after 24h incubation at [amphiphile] = 160 μM in full DMEM (10% FBS). Total fluorescence (first row) or ratiometric images of unimer/micelle pixel ratio after background removal (second row) are shown in the basal lamina compartment (A) or at the edge of an MCF7 spheroid (C). Scale bars represents 20 μm. Intensity profiles (B) of 3×80 μm rectangles inside the basal lamina compartment are shown for total fluorescence images (blue line) and ratiometric images (pink line). The mean fluorescence intensity or unimer/micelle ratio is quantified for each chip compartment (D): flow channel (control), basal lamina, tumor ECM (outside spheroids) and inside spheroids (N≥3). A horizontal gray line is drawn for visualization purposes, corresponding to a unimer/micelle ratio if 1.

In order to differentiate if the interaction that we observed is happening more in the micellar or unimer forms, we checked the ratiometric images (Figure 4A bottom row). The PEtOx-Non and PEG-Non showed uniform micelle conformation (indicated by green color), which could be expected based on being relatively more stable. On the other hand, PAA polymers had an overall higher unimer/micelle ratio, of ∼1.5 for PAA-Non and ∼2 for PAA-Hex (Figure 4B). In this case, some disassembly is already happening in solution, probably due to their higher tendency for interaction with serum proteins.^19^ The ratio profile in the basal lamina appears more noisy for the PAA micelles/unimers, but without a clear difference in the regions corresponding to bright structures in the total fluorescence image. Thus, we concluded that PAA polymers are binding to the basal lamina in a similar equilibrium state as in media solution (predominantly in unimer form) and that the ECM binding caused further destabilization as the unimer/micelle ratios are higher than the ones in the control media (Fig 4D). As for the PEtOx-Hex and PEG-Hex, ratiometric images showed faint traces of increased unimer signal, causing a slightly higher mean ratio in basal lamina compartment compared to the ratio in solution (Figure 4D). Since micelles of PEtOx-Hex and PEG-Hex are more stable than PAA micelles, but less stable than their Nonyl analogues, we can assume that the free unimers in solution are more likely to bind to ECM, while most micelles remain in a stable form in solution. Overall, basal lamina binding was highly influenced by micelle surface charge, with PAA polymers showing the most binding, and also by micelle stability, with PetOx-Hex and PEG-Hex interacting more than their Nonyl counterparts.

In the “tumor ECM” compartment of the chip, MCF7 spheroids were embedded in a mix of collagen type I and hyaluronic acid. Unlike the basal lamina compartment, our “tumor ECM” gel had no visible impact on the total fluorescence signal nor on the micelle distribution, except for PAA polymers which showed an increase in fluorescence, but nearly half compared to the increase observed in the basal lamina compartment (Figure 4D). In this case, the PAA micelles might be already destabilized after their passage through basal lamina. Interestingly, when quantifying the mean unimer/micelle ratio in the “tumor ECM”, all Hexyl polymers showed higher ratios (similar to the ones in basal lamina) while Nonyl polymers maintained a ratio close to control. This clearly points to the stabilizing effect of the longer hydrophobic tails leading to less ECM interactions. Overall, the “tumor ECM” gel had little impact on micelle passage. We found differences however in the uptake behavior into MCF7 spheroids. Using the total fluorescence images, we quantified the mean intensity inside spheroids as an indicator of cellular internalization. Overall, we observed that PEtOx-Hex and PEG-Hex had similar uptake. The more stable PEtOx-Non and PEG-Non showed weaker intensities inside the spheroids, indicating on their lower degree of internalization, while PAA micelles had the highest intensity. This is consistent with previous uptake experiments in 2D HeLa cell cultures,^19^ in which we observed differences in intracellular distribution, with PAA formulations showing membrane signal, while the others were internalized in endocytic vesicles. The 3D setting inside the chip posed imaging limitations (due to sample thickness, and the gel and spheroid densities), which did not allow a clear assessment of the intra-cellular distribution, but it seemed to follow the same trend.

Ratiometric images of the spheroids indicated predominantly the unimer form inside cells for all polymers, with unimer/micelle ratios above 3. Having only unimers inside the cells is expected for the long incubation time used with the chip (24h). It is not excluded that polymers internalize in micelle form and break down inside the cells, but this process would be fast and difficult to capture in the given conditions. For PAA containing polymers, the ratio was higher also outside cells due to their lower stability during a long incubation time in full DMEM. As we showed previously with serum albumin experiments, PAA micelles tend to dissociate in solution due to protein interaction.^19^

### FRAP

Next, we assessed unimer and micelle dynamics in different chip compartments using fluorescence recovery after photobleaching (FRAP). Briefly, a circular bleach region of 35.3 μm was exposed to high intensity 405 nm laser, causing local photo-bleaching of the coumarin labels. The diffusion of amphiphiles outside of the bleached area, being replaced by ones with intact fluorescence from the surroundings, causes a local recovery of fluorescence signal. The signal was recorded using split unimer/micelle channels as explained in the confocal imaging section, in order to obtain the diffusion constants of both unimers and micelles in different chip compartments.

Firstly, looking at the normalized recovery curves we can say that all of them had a complete recovery – we did not observe an immobile fraction (Figure 5, Supplementary figure 3). This is indicative of the nature of the possible binding, meaning that any occurring interactions were not strong enough to immobilize the bleached molecules for the duration of the FRAP acquisition. Instead, the exchange of bright and dark amphiphiles happened rather fast.

**Figure 5.**
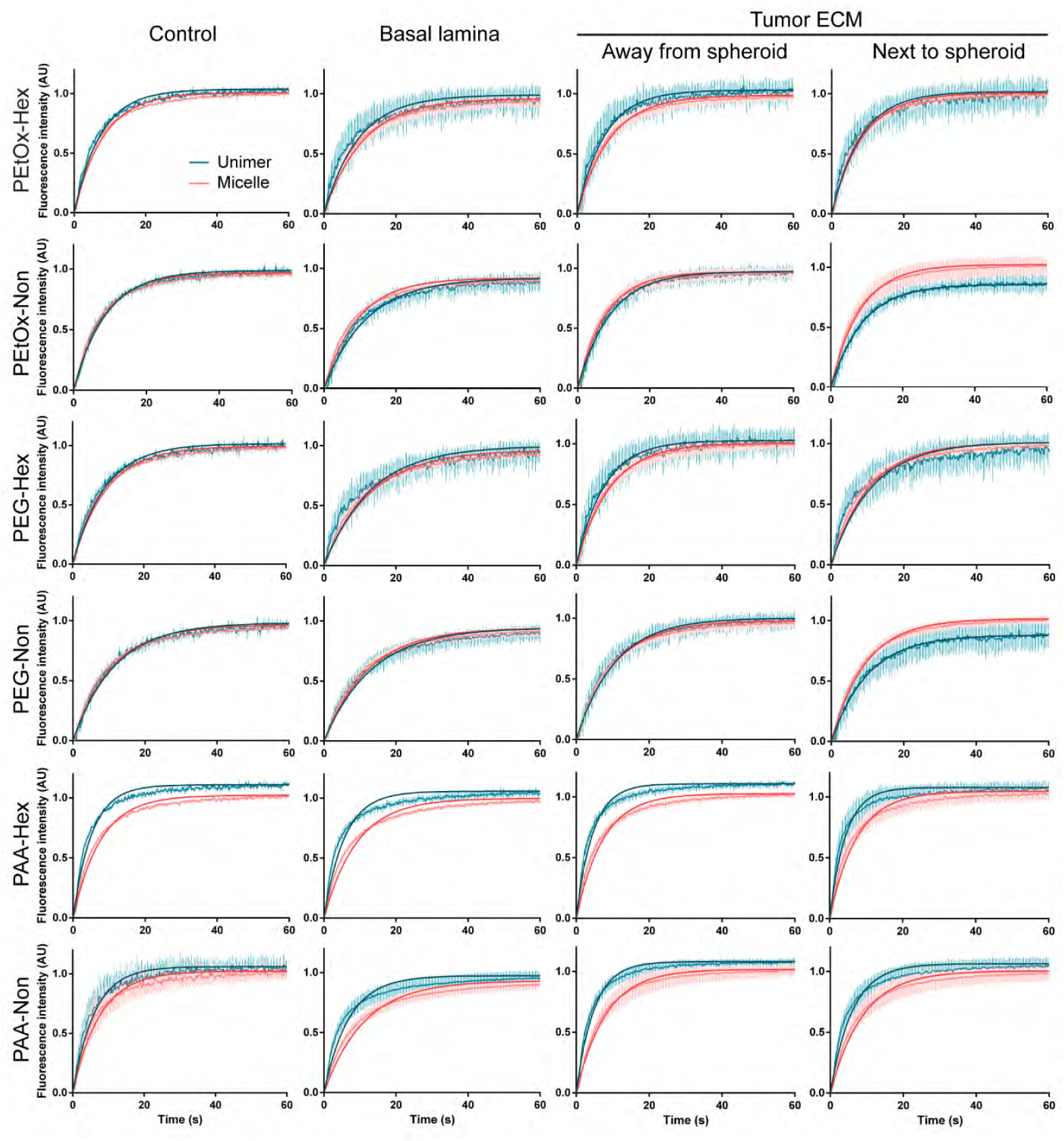
FRAP recovery curves of unimer (blue) and micelle fluorescence (red) represented as mean and standard deviation (faint lines), with fitted one-component exponential curve (dark lines), shown in different parts of the chip. Control measurements represent micelles in solution (in full DMEM, 10% FBS).

Secondly, we observed a slightly slower recovery rate inside the basal lamina for all formulations, which is reflected in lower diffusion constants (Figure 6). Being present in all formulations, we can assume it to be due to geometric hindrance imposed by the microarchitecture of the basal lamina mesh inside the bleach area. However, the Hexyl amphiphiles seem to be affected more than the Nonyl ones and the difference was higher for PAA compared to PetOx and PEG. In these cases we can assume the slower diffusion is caused by interactions with basal lamina structures.

**Figure 6.**
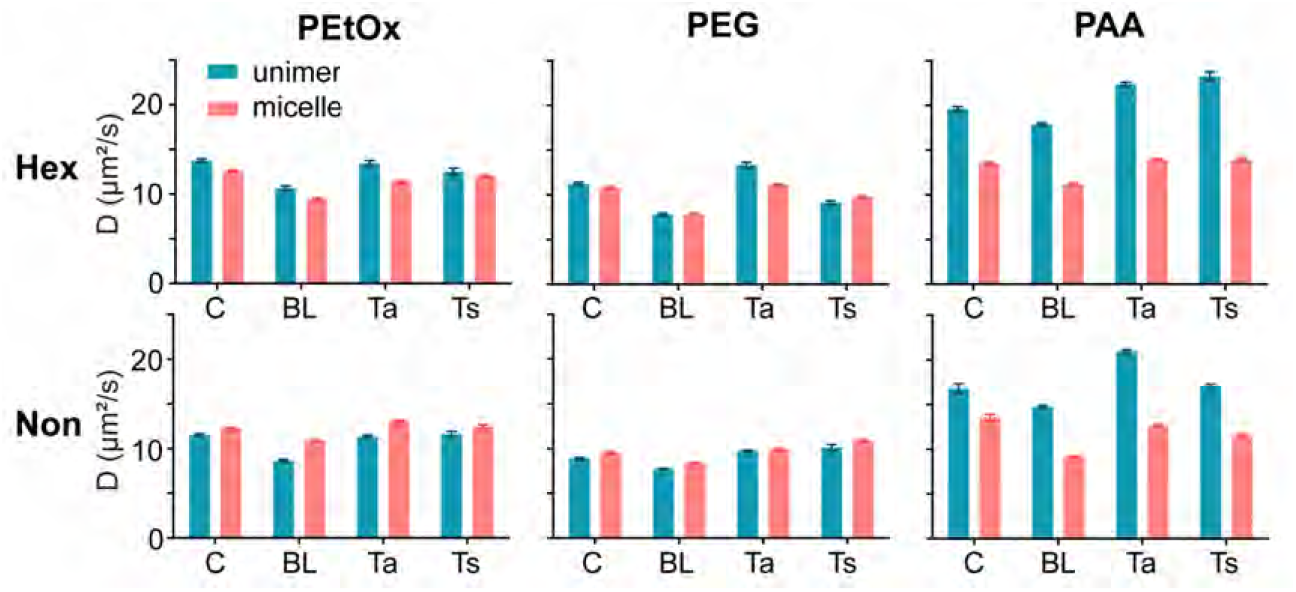
Diffusion constants of unimers (blue) and micelles (red) calculated from FRAP measurements in different locations inside the chip. Control measurements (“C”) were performed in solution, on a glass slide. “BL”, “Ta”, “Ts” represent basal lamina, tumor ECM away from spheroids and tumor ECM next to spheroids respectively.

In the “tumor ECM”, the diffusion is similar to the one in solution, indicating that collagen type I and hyaluronic acid posed no hindrance to micelle diffusion. This is in accordance with the results of confocal imaging, where we observed little interaction with the “tumor ECM”.

The FRAP data should be seen as reflecting the overall interactions rather than strictly the diffusion rate. By looking at the difference in size between a free unimer (expected to be 5-10 nm) and a formed micelle (15-25 nm in diameter), one might expect to observe a difference in diffusion rate in the FRAP data. However, one challenging aspect in interpreting the FRAP data is the presence of different types of interactions in our system. One of these interactions is the dynamic equilibrium of unimers and micelles. The transition between free unimers and the ones assembled into micelles is probably happening very fast, meaning that although we measure separately the unimers and micelles fluorescence, there are probably unimers that get back into micelle form and vice-versa during the FRAP acquisition. Another factor is the presence of serum proteins (experiments are performed in full DMEM media, with 10% FBS) which is likely influencing the measured diffusion rate. In our previous study, we showed that serum albumin is binding to both unimers and micelles causing destabilization.^19,27^ Thus, protein-bound unimers are likely more bulky and diffusing slower than a free unimer. This means the diffusion constant of a protein-bound unimer can be closer to the one of a micelle with less protein interaction, which is the case for the more stable PEtOx and PEG formulations. For PAA, the charged shell causes a higher interaction with proteins, which can affect both unimers and micelles, being reflected in a clear difference of unimer to micelle diffusion constants.

We measured the diffusion in “tumor ECM” close to and away from spheroids in an attempt to check if our MCF7 spheroids are affecting the ECM diffusion, by either stiffening or degrading the matrix in their proximity. However, we did not observe a difference between these locations. Other studies have shown ECM remodeling by tumor cell spheroids,^29^ as well as stiffening and hindered diffusion due to collagen deposition by fibroblasts.^30^ In this sense, we can conclude that our model was too simple to measure this difference. It could be the relative short time in the chip (24h) or the spheroids being too small (due to limitations given by channel size) to have a visible impact on the ECM conformation. Probably a different cell type or a model containing co-cultures of cancer and stromal cells would be able to recapitulate these features, although with added degrees of complexity. Alternatively, nanoparticle diffusion has been studied in the ECM deposited intercellularly inside spheroids,^31^ which could be an interesting approach for future studies.

Our study reveals a positive correlation between ECM binding of polymeric micelles and their uptake by spheroid cellular, with charged polymers (PAA) showing reversible binding to ECM and higher uptake in MCF7 spheroids. A similar correlation was found by Valente et al. for 10 nm gold nanoparticles of different surface charges,^32^ highlighting the importance of testing both ECM penetration and cellular uptake in relevant 3D models.

## Conclusions

In summary, the current article presents a simple tumor-on-a-chip model to mimic the tumor tissue barrier, consisting of two types of ECM (basal lamina and collagen type I – hyaluronic acid mix) and MCF7 spheroids. Inside this 3D chip model we tested the distribution and mobility of polymeric micelles, comparing three hydrophilic shells: PEtOx, PEG and PAA, with two lengths of the hydrophobic ends (Hexyl or Nonyl). We observed different interaction behaviors inside the basal lamina, correlated with micelle stability: avoidance of basal lamina mesh for the more stable PEtOx-Non and PEG-Non and reversible binding for negatively charged PAA formulations. Spheroid uptake, on the contrary, was best for PAA formulations, emphasizing the importance of testing both cellular uptake and delivery through ECM. Overall, the study showcases the use of a simple microfluidic chip for testing the interactions of polymeric nanocarriers with biological interfaces and the potential of such simple test models to significantly contribute towards increasing the understanding of tumor drug delivery systems in a much shorter and efficient feedback loop between formulation and testing.

## ASSOCIATED CONTENT

### Supporting Information

Cytotoxicity assay, additional confocal images and FRAP recovery curves. This material is available free of charge.

## AUTHOR INFORMATION

### Notes

The authors declare the following competing financial interest(s): R.H. and V.R.D.L.R. are co-founders of Avroxa BVBA that commercializes poly(2-oxazoline)s as Ultroxa. The other authors have no conflicts of interest to declare.

### Author Contributions

‡A.R.O. and A.J. authors contributed equally.

## ACKNOWLEDGMENTS

This project has received funding from the European Union’s Horizon 2020 research and innovation program under the Marie Skłodowska-Curie [grant agreement no. 765497 (THERACAT)]. R.J.A thanks the ISRAEL SCIENCE FOUNDATION (grant No. 1553/18) for the support of this research. S.T. thanks the ADAMA Center for Novel Delivery Systems in Crop Protection, Tel-Aviv University for the financial support. G.S. thanks the Marian Gertner Institute for Medical Nanosystems in Tel Aviv University for their financial support. R.H. and V.R.D.L.R are grateful to Ghent University and FWO for continuous financial support. S.P. acknowledges the financial support by the Spanish Ministry of Science and Innovation (PID2019-109450RB-I00/AEI/10.13039/501100011033).

## ABBREVIATIONS

7-DEAC: 7(diethylamino)coumarin-3-carboxylic acid;
DDS: drug delivery system;
DMEM: Dulbecco’s Modified Eagle Medium;
FBS: fetal bovine serum;
FRAP: fluorescence recovery after photobleaching;
HA: hyaluronic acid;
ECM: extracellular matrix;
PAA: poly(acrylic acid);
PBS: phosphate buffered salide;
PEG: poly(ethylene glycol);
PEtOx: poly(2-ethyl-2-oxazoline).

