## Supplementary figures for "Reaching the tumor: mobility of polymeric micelles inside an *in vitro* tumor-on-a-chip model with dual ECM"

### **Supplementary information**

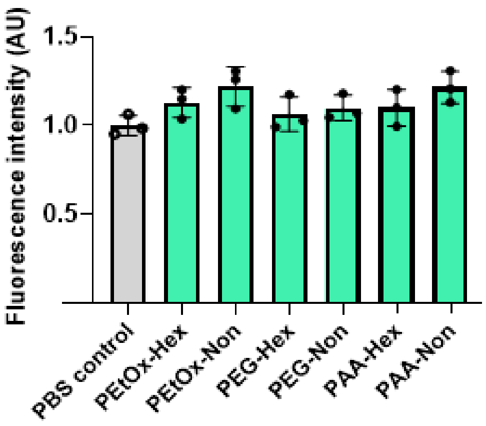


Supplementary figure 1: Cytotoxicity assay of micelles on MCF7 cells, assessed with Presto Blue.

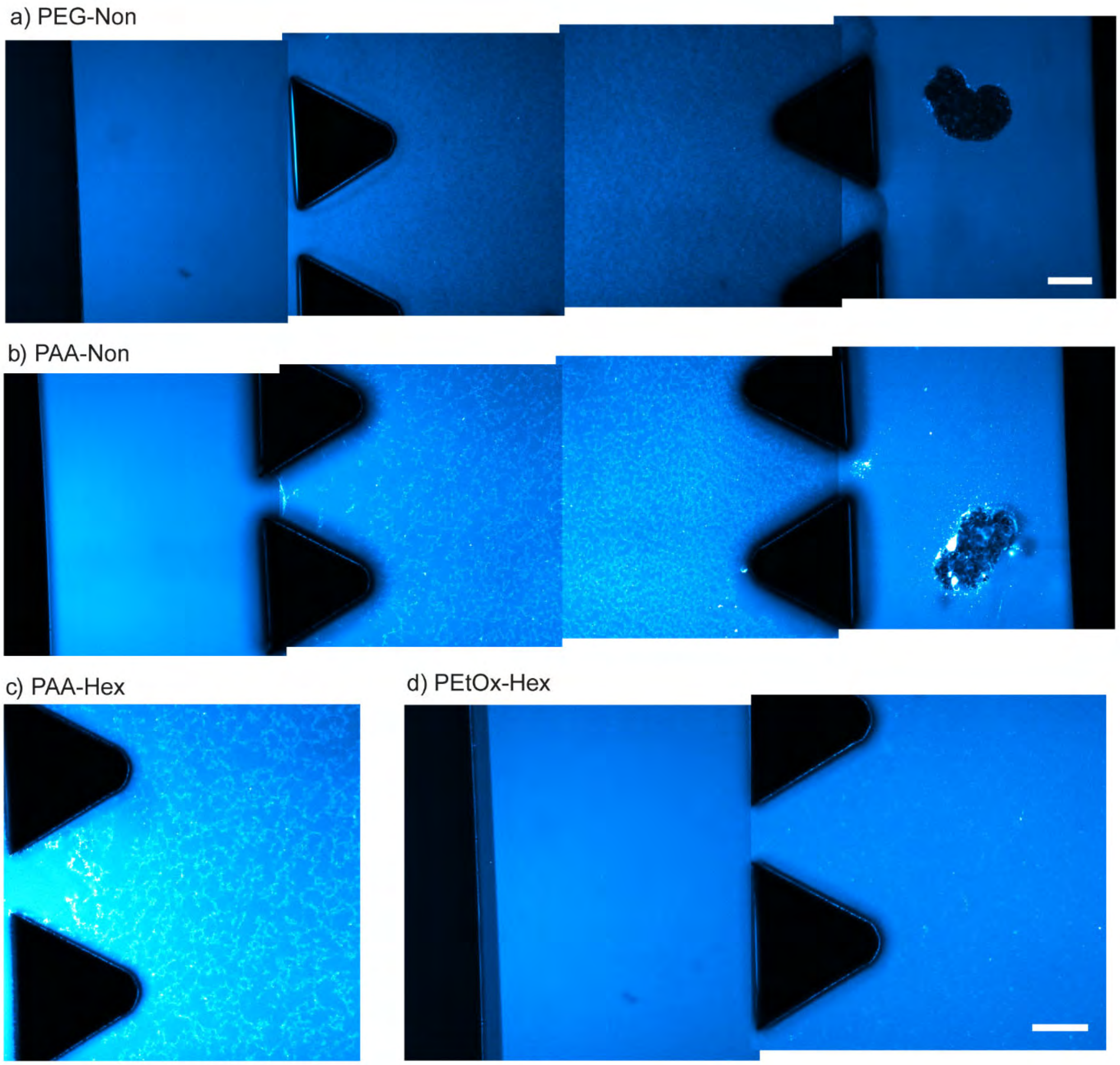


Supplementary figure 2: Overview images (total fluorescence) of coumarin-labeled micelles inside multigel chip model. The chip contains three channels separated by triangular pillars; the left channel contains only solution of micelles in full DMEM media, the middle channel contains a basal lamina gel model and the right channel contains MCF7 spheroids embedded in a mix of collagen type I and hyaluronic acid. The gel structure inside the middle channel was visible after micelle adition in some cases, either as a "dark" structure for more hydrophobic micelles (a – PEG-Non), or as a bright structure for PAA micelles (b – PAA-Non, c – PAA-Hex). In-between, the less hydrophobic micelles did not reveal the structure, but only a few bright spots when compared to the left channel (d – PEtOx-Hex). Scale bars represent 100 µm.

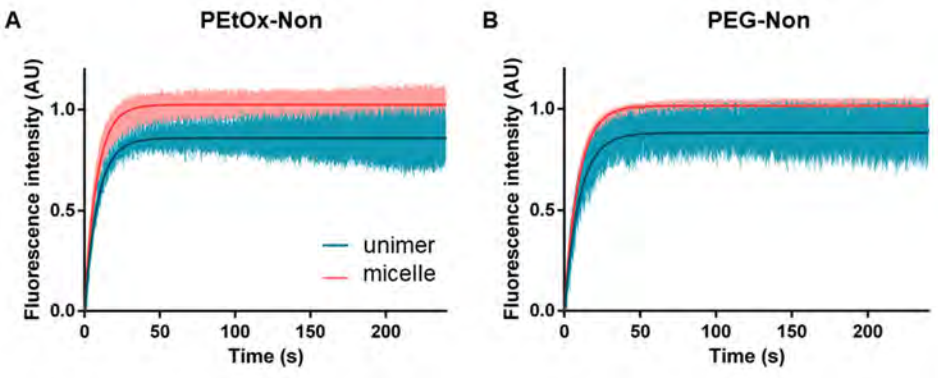


Supplementary figure 3: Zoom-out of FRAP recovery curves for PEtOx-Non (A) and PEG-Non (B) in collagen-HA next to spheroid, showing the mean fluorescence in the bleach area during the entire post-bleach acquisition time. Graph plotted as mean +/-SD. Unimer signal is shown in blue, while micelle signal is shown in red.

### Supplementary methods

#### Cytotoxicity in 2D culture of MCF7 cells

MCF7 cells were seeded in a flat-bottom transparent 96-well plate (Nunc, Thermo Scientific) as 5000 cells/well and allowed to grow for 24h. The supernatant was replaced by 160µM micelle solution in full DMEM (10% FBS). PBS (pH 7.4) or Triton X-100 0.01% v/v in full DMEM were used as negative or positive controls respectively. After 24h, the cells were incubated for 1h with Presto Blue solution (ThermoFisher) as 10% v/v, at 37° C, 5% $CO_2$ . Florescence was measured using a multimode microplate reader (Infinite M200 Pro, Tecan), with 550 nm excitation, acquiring the signal at well bottom at 600 nm emission. Samples were prepared in triplicate, in randomized order.
